# CreC Sensor Kinase Activation Enhances Growth of *Escherichia coli* in the Presence of Cephalosporins and Carbapenems

**DOI:** 10.1101/615310

**Authors:** Yuiko Takebayashi, Emma S. Taylor, Sam Whitwam, Matthew B. Avison

## Abstract

Mutants with enhanced growth in the presence of antibiotics but where the antibiotic’s minimum inhibitory concentration does not change, are difficult to identify in mixed populations because they are not amenable to selection. Here we report that activatory mutations in the CreC signal sensor enhance growth of *Escherichia coli* in the presence of cefoxitin, cefotaxime and meropenem without increasing their minimum inhibitory concentrations. Enhanced growth is dependent on overproduction of the inner-membrane cre-regulon protein CreD.

## Text

In *Escherichia coli*, CreC is a sensor kinase and CreB is a response regulator and together they form a two-component regulatory system that controls the expression of the cre regulon, a group of genes with poorly characterised functions (1–3). We have previously shown that an activatory mutation in CreC confers the Cet phenotype: tolerance of the protein antibiotic Colicin E2, through over-production of the protein YieI (3). Whilst the mechanism by which YieI confers colicin resistance is not clear it may be due to modifications in the outer envelope which restrict entry of the colicin to the cell (3).

To learn more about the Cet phenotype, we used phenotype microarray analysis to characterise differences between CTX6, a well-characterised Cet mutant (3), versus CTX6Δ*creB*, where cre regulon hyper-expression is ablated and the Cet phenotype is reversed (3). Phenotype microarray analyses were performed by BIolog (Hayward CA, USA). Of almost 2000 growth conditions tested in the phenotype microarray, significantly improved growth of CTX6 versus CTX6Δ*creB* was seen in media with 13 different chemical additions. Eight of these chemicals are β-lactam antibiotics (Table 1). MICs of β-lactams were not noticeably higher against CTX6 than CTX6Δ*creB*, according to E-test and broth microdilution methodologies (data not shown) but the CreC activatory mutation in CTX6 was reproducibly seen to enhance growth at sub-MIC concentrations of some β-lactam drugs during growth curve analysis. For these assays, 500 µl of an overnight culture grown in LB medium (Oxoid) were used to inoculate 10 ml of fresh LB medium in a sealed universal bottle and the antimicrobial drug of interest was added. The starting optical density at 600 nm (OD_600_) of each subculture was approximately 0.1. Cultures were incubated at 37°C with 150 rpm shaking and O.D._600_ measured every hour using a spectrophotometer by taking 1 ml from the culture. For example, *E. coli* MG1655 (the parent of CTX6) suffers an initial lag in growth in the presence of half the MIC of cefoxitin (Fig 1A) or cefotaxime (Fig 1B) (both from Sigma). Later, a post-antibiotic effect is seen, and growth of the population starts up again (Fig. 1). CTX6 does not suffer such an obvious lag in growth and there is a significant enhancement in optical density at 180 or 360 min after the addition of cefoxitin or cefotaxime, respectively (*p* <0.05 for both). CTX6Δ*creB* displays the wild-type phenotype (Fig. 1) confirming that CreBC hyper-activation is responsible for enhanced growth in the presence of these β-lactam antibiotics.

**Table 1.** Chemicals in Phenotype Microarray analysis where significant changes in growth – measured as an increased area under the growth curve (AUC) – were seen in CTX6 (Cet) versus CTX6Δ*creB*.

| **Moderate Effect**<br>**(26-75% increase in AUC)** | **Major Effect**<br>**(>75% increase in AUC)** |
| --- | --- |
| Nafcillin | Amoxicillin |
| Ampicillin | Cefotaxime |
| Cefuroxime | Aztreonam |
| Cefoxitin | Phenylarsine oxide |
| Carbenicillin |  |
| Cetylpyridinium chloride |  |
| Thallium Acetate |  |
| 9-Aminoacridine |  |
| Sodium Arsenate |  |

**Figure 1.**
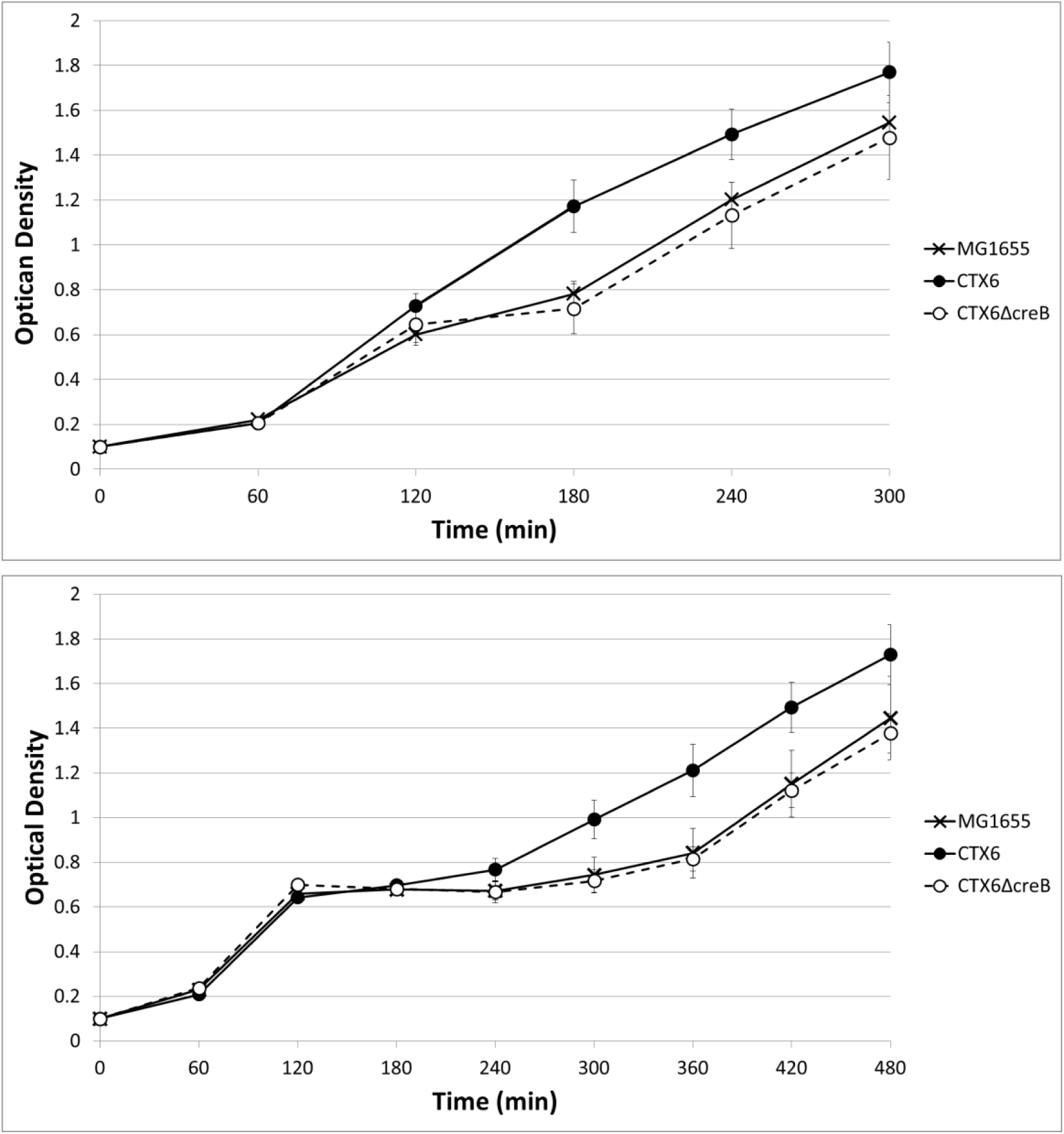
Effect of Cet phenotype on *E. coli* growth curve in the presence of cefoxitin or cefotaxime. Optical density of an LB culture was measured at 600 nm every hour following subculture and addition of antibiotic (time zero): top graph, cefoxitin; bottom graph, cefotaxime, each used at half its MIC against *E. coli* MG1655. Data are mean plus/minus standard error of the mean, n=6

Microarray transcriptomics has previously revealed that six genes are upregulated >10-fold in CTX6 in a CreB-dependent manner (3). The use of previously constructed deletion mutants of these genes in CTX6 (3) revealed that *creD* is the CreBC regulated gene responsible for enhanced growth of CTX6 in the presence of cefoxitin, as illustrated in figure 2. Deletion of *creD* in the CTX6 background reduced the optical density of a culture at 180 min post addition of the drug to the same extent as deletion of *creB* or *creC* (p<0.005 for each).

**Figure 2.**
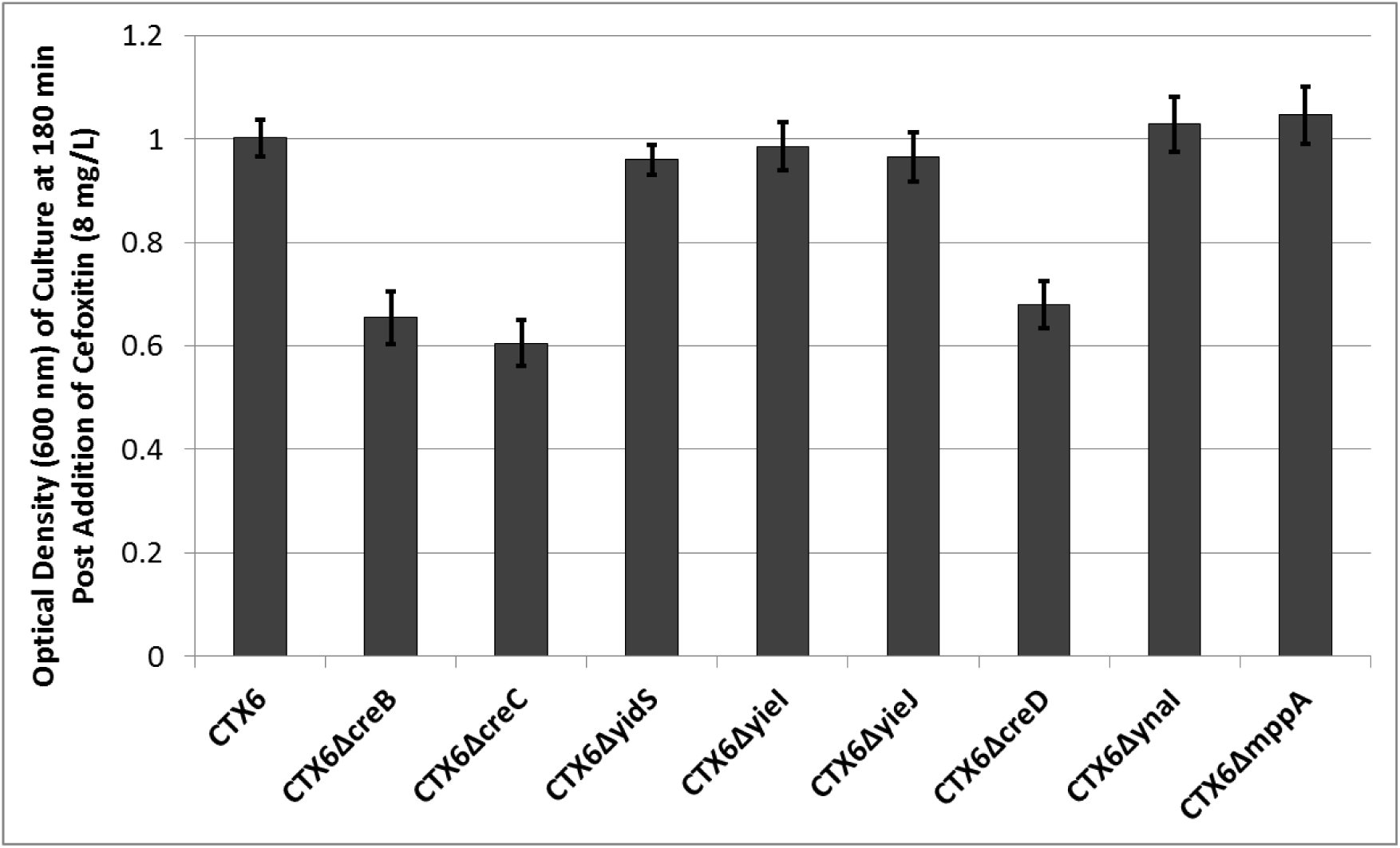
Effect of deletion of various Cre regulon genes in the *E. coli* Cet mutant CTX6 on growth in the presence of cefoxitin. Optical density of an LB culture was measured at 600 nm every hour following subculture and addition of antibiotic (time zero). Data represent means plus/minus standard error of the mean, n=3 after 180 mins post incubation.

It is possible to activate cre regulon gene expression by over-expressing the response regulator *creB* in an otherwise wild-type background strain. We did this, as previously, using an arabinose expression system (2) and found that it enhances growth in the presence of β-lactam antibiotics, even at antibiotic concentrations greater than the MIC. For example, with cefotaxime, the growth curve of MG1655 carrying a control plasmid shows the drug inhibiting growth, lysing cells and overwhelming the population, causing the optical density of the culture to reduce to basal levels (Fig. 3). Carriage of a plasmid allowing *creB* over-expression does not affect the MIC – the drug still kills the population – but the growth curve is more pronounced. There is more growth prior to killing, as seen particularly at 120 min post cefotaxime treatment in figure 3 (p<0.01). Deletion of the CreBC regulated gene *creD* blocks the ability of CreB hyper-production to improve the growth of MG1655 (p<0.05 at 120 min post cefotaxime treatment [Fig 3]) confirming a role for CreD over-production in this CreBC mediated β-lactam tolerance phenotype.

**Figure 3.**
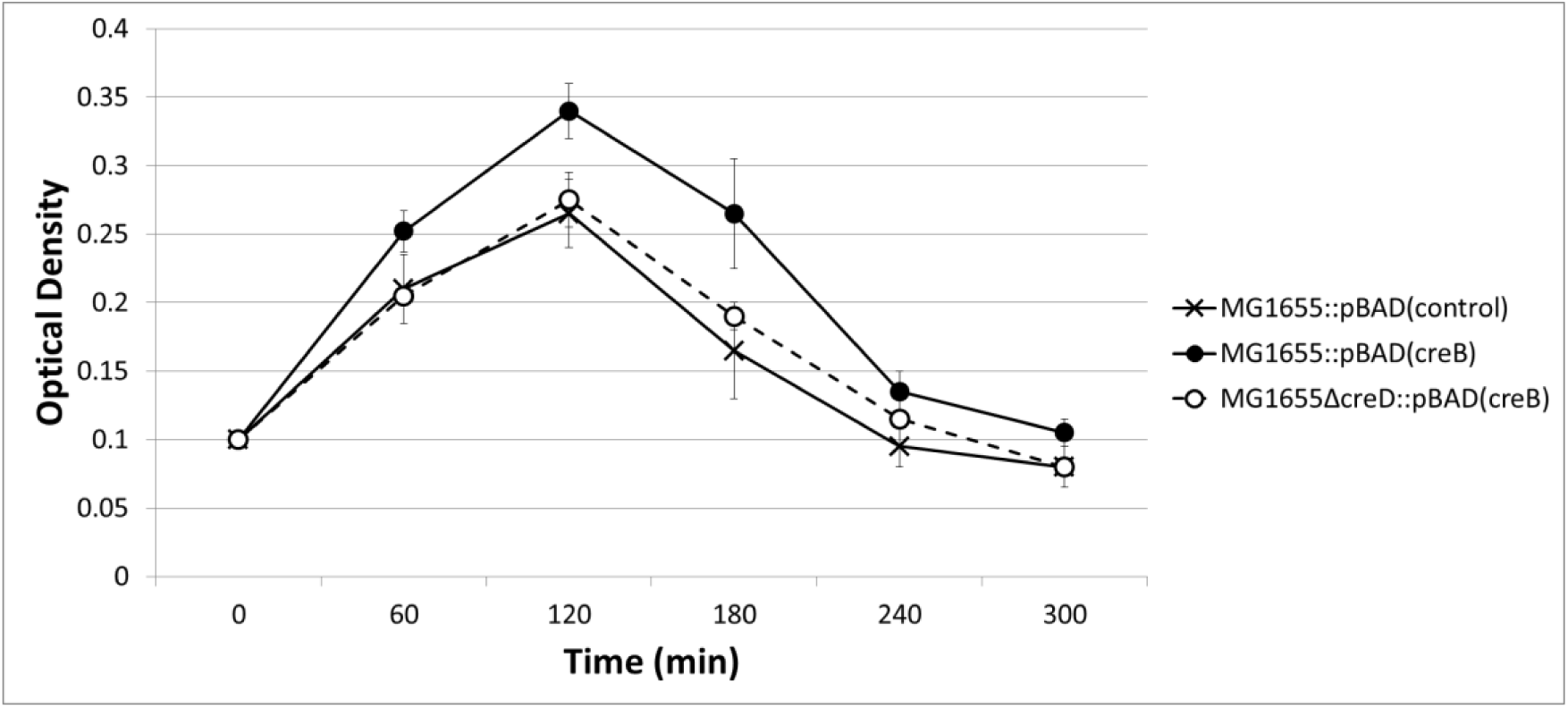
Effect of activating the Cre regulon through over-production of CreB on *E. coli* growth in the presence of cefotaxime. Optical density of an LB culture was measured at 600 nm every hour following subculture and addition of cefotaxime (time zero) used at half its MIC against *E. coli* MG1655. Arabinose (0.2 % w/v) was added in all growth media to stimulate CreB production from the pUB6073 (pBAD(*creB*)) plasmid as used in (2). Data represent means plus/minus standard error of the mean, n=3 after 180 mins post incubation.

To see if the *creC* hyperactive mutation in CTX6 gives a fitness advantage relative to CTX6Δ*creB* over repeated cycles of growth, pairwise competition experiments were performed over 4 days. In order to mark CTX6Δ*creB* so that it could be quantified in a culture when mixed with CTX6, the derivative CTX6*creB*::Chl^R^ was used. Here, *creB* has been disrupted by insertion of a chloramphenicol resistance gene rather than deleted as used in earlier experiments, though the effect on CreBC regulated gene expression is known to be identical (2). CTX6*yieJ*::Chl^R^ (3) was used in parallel as a control to confirm that any change in fitness seen was specific to disruption of *creB* and not due to Chl^R^; *yieJ* is not responsible for the growth enhancement phenotype of CTX6 (Fig. 2).

To perform these experiments, initially, cultures of both strains to be competed were inoculated separately into LB and incubated for 24 h at 37°C with shaking at 160 rpm. Next, 5 µl of each culture was used to inoculate a separate flask containing 50 ml of DM25 minimal medium (4), which was prepared from Davis minimal medium (Difco, Oxford, UK) supplemented with glucose and thymine (25 mg.l^−1^ and 2 mg.l^−1^ respectively). Cultures were incubated at 37°C with shaking as above. After overnight incubation, 500 µl of each culture was transferred into a fresh 50 ml aliquot of DM25 minimal medium, again separately, and the inoculated medium was divided into six screw top universal bottles and incubated at 37°C overnight with shaking, as above. Upon inoculation, these cultures were referred to as day zero cultures. Day one (mixed) culture started with 250 µl each of the two day zero cultures to be competed being mixed together in the same flask containing 50 ml of fresh DM25 medium containing a β-lactam antibiotic as necessary (i.e. there were six flasks for each competition experiment). The day one (mixed) cultures were incubated for 24 h at 37°C with shaking. Each day one culture was then sub-cultured by transferring a 500 µl of the culture into a separate flask containing 50 ml of DM25 medium (containing the same β-lactam, if necessary) to generate day two (mixed) cultures. The last step was repeated daily until four days of mixed cultures had passed. The competition between the two strains was measured by performing a serial dilution and counting the number of colony forming units (cfu) per ml of CTX6 and of CTX6::Chl^R^ in each mixture at the end of each day (including day zero). CTX6::Chl^R^ was counted following plating on LB agar containing 30 mg.l^−1^ chloramphenicol. CTX6 was counted by plating on LB agar with no antibiotic and subtracting the CTX6::Chl^R^ count. The selection rate constant (r) was used as a measure to estimate the fitness cost of Chl^R^ insertion after each day of the competition by comparing M, the Malthusian parameter for each strain in the competition (5) so that M = ln(N_1_/N_0_), where N_0_ is the density of the strain (cfu/ml) at the start of the day (density at the end of the previous day divided by 100 to take account of the dilution factor on subculture) and N_1_ is the density of the strain (cfu/ml) at the end of the day. The selection rate for a competition is therefore calculated as r = M_1_-M_2_, Where M_1_ relates to CTX6 and M_2_ relates to CTX6::Chl^R^. For each competition (one mixed culture) there were four selection rate values, one for each day, and for each fitness cost experiment, 6 competitions were run. Hence for each competition between two strains, 24 r value datapoints are obtained. Differences in these sets of r value data for different comparisons were assessed using an unpaired t-test with Welch’s correction to assess the statistical significance of the differences observed.

In the absence of antibiotics, disruption of neither *creB* nor *yieJ* had any significant effect on the fitness of CTX6 (Table 2). However, in the presence of half the MICs of cefotaxime, cefoxitin or meropenem (AdooQ Bioscience), an approximately 15% per day reduction in relative fitness (W) was observed after disrupting *creB* (p<0.05 for all comparisons) though there was no significant effect of disrupting *yieJ* (Table 2). We also performed competition experiments using ampicillin or ceftazidime (both from Sigma), but no significant effect of CreBC hyperactivation on fitness was seen (Table 2). To put the observed changes in relative fitness into perspective: starting with a ratio of approximately 1:1 for the two competing strains, after 4 days of competition the ratio was approximately 10:1 in favour of the CreBC hyperactive strain in the presence of half the MIC of cefoxitin, cefotaxime or meropenem.

**Table 2.** Effect on fitness of disrupting *creB* in CTX6 measured in pairwise comparison with CTX6 during growth in the presence of half MIC of various antibiotics. Fitness was measured over four days, and is an average of six repetitions, as set out in the text. Data are mean percent fitness difference per day. Negative values mean that the ChlR derivative is less fit than CTX6 in pairwise competition. Stars represent differences where p<0.05 according to an unpaired t-test with Welch’s correction.

| Competition | Relative Percent Fitness of *Chl<sup>R</sup>* strain |
| --- | --- |
| CTX6*creB*:*Chl<sup>R</sup>* vs CTX6<br>(No Antibiotic) | +3.0 +/- 3.4 |
| CTX6*yieJ*:*Chl<sup>R</sup>* vs CTX6<br>(No Antibiotic) | -4.9 +/- 3.9 |
| CTX6*creB*:*Chl<sup>R</sup>* vs CTX6<br>(Plus Ampicillin) | +4.4 +/- 2.8 |
| CTX6*creB*:*Chl<sup>R</sup>* vs CTX6<br>(Plus Ceftazidime) | -0.7 +/- 4.8 |
| CTX6*creB*:*Chl<sup>R</sup>* vs CTX6<br>(Plus Cefoxitin) | -17.7 +/- 3.4* |
| CTX6*yieJ*:*Chl<sup>R</sup>* vs CTX6<br>(Plus Cefoxitin) | -2.0 +/- 4.7 |
| CTX6*creB*:*Chl<sup>R</sup>* vs CTX6<br>(Plus Cefotaxime) | -12.4 +/- 4.9* |
| CTX6*creB*:*Chl<sup>R</sup>* vs CTX6<br>(Plus Meropenem) | -15.9 +/- 4.1* |

## Conclusions

Antibiotic resistance is defined in terms of a minimum inhibitory concentration (MIC) of an antibiotic measured following growth of a very dilute starting culture in the presence of that antibiotic. After a set period of incubation, the MIC is defined as the concentration of drug required to completely inhibit growth. If the MIC of a drug against a bacterial isolate is greater than a consensually defined breakpoint – based on clinical experience and pharmacokinentic/dynamic properties of the antibiotic – the isolate is said to be resistant (6). However, in vivo, antibiotic concentrations are dynamic, and there are periods when they are sub-MIC, so even susceptible bacteria can multiply before the drug concentration raises to overcome it. Whilst this does not, of itself, cause treatment failure, it might provide time for resistant mutants to emerge, or for symptoms to persist in the patient before a cure is established. Because of this possibility, we are interested in mutations that confer growth enhancement in the presence of an antibiotic without necessarily altering the antibiotics’ MIC. Such mutations are very difficult to identify in a population because, in contrast to resistant mutants, enhanced growth mutants are not amenable to easy selection in vitro. Accordingly, even if the growth-enhancing mutation were a high-frequency event, the proportion of a population displaying the growth enhancement phenotype would be so small it would essentially be invisible. This phenotype of enhanced growth in the presence of antibiotic is somewhat different from the much-discussed phenotype of “persistence”, where bacteria reduce their metabolic activity and/or growth rate to escape from the lethal actions of antibiotics, which tend to only kill growing cells (7). The persister phenotype is easier to select than the growth enhancement phenotype because the persister population reveals itself after removal of antibiotic (8); the enhanced growth phenotype population does not.

In this paper, we have identified a mechanism that confers growth enhancement in the presence of β-lactams, the most commonly prescribed antibiotics, in *E. coli*, one of the most commonly encountered human pathogens. The clinical relevance of this finding is unclear, but it is an interesting observation that activation of the production of CreD, an inner membrane protein of uncertain function, confers growth enhancement in the presence of β-lactams. Interestingly, CreBC shares significant sequence identity with the BlrAB two-component system in *Aeromonas* spp (9,10), which is activated in response to β-lactam challenge, and controls β-lactamase production and so confers β-lactam resistance (11). Importantly, BlrAB also activates transcription of *blrD*, which is a homologue of *E. coli creD* (10). Indeed, a similar two component system, CreBC/BlrAB in *Pseudomonas aeruginosa* is also activated during β-lactam challenge (12,13), and whilst it does not control β-lactamase production, it does control transcription of the *creD* homologue in this species (12,13). It is therefore striking, that CreBC activation in *E. coli* enhances growth in the presence of β-lactams. This speaks to a common function for the CreD/BlrD proteins in these varied species of gamma-proteobacteria.

## Acknowledgements

We thank Dr Christopher D. Smith for preliminary analysis of Cet mutants and Dr James L. Cariss for assisting with the phenotype microarray work.

## References

1. Avison MB, Horton RE, Walsh TR, Bennett PM. (2001). Escherichia coli CreBC is a global regulator of gene expression that responds to growth in minimal media. J Biol Chem. 276:26955–61.

2. Cariss SJ, Tayler AE, Avison MB. Defining the growth conditions and promoter-proximal DNA sequences required for activation of gene expression by CreBC in Escherichia coli. J Bacteriol. 190:3930–9.

3. Cariss SJ, Constantinidou C, Patel MD, Takebayashi Y, Hobman JL, Penn CW, Avison MB. (2010). YieJ (CbrC) mediates CreBC-dependent colicin E2 tolerance in *Escherichia coli*. J Bacteriol. 192:3329–36.

4. Enne VI, Bennett PM, Livermore DM, Hall LM. (2004). Enhancement of host fitness by the sul2-coding plasmid p9123 in the absence of selective pressure. J Antimicrob Chemother. 53:958–63.

5. Travisano M, Lenski RE. (1996). Long-term experimental evolution in *Escherichia coli*. IV. Targets of selection and the specificity of adaptation. Genetics. 143:15–26.

6. Mouton JW, Brown DF, Apfalter P, Cantón R, Giske CG, Ivanova M, MacGowan AP, Rodloff A, Soussy CJ, Steinbakk M, Kahlmeter G. (2012). The role of pharmacokinetics/pharmacodynamics in setting clinical MIC breakpoints: the EUCAST approach. Clin Microbiol Infect. 18:E37–45.

7. Lewis K. (2010). Persister cells. Annu Rev Microbiol. 64:357–72.

8. Rowe SE, Conlon BP, Keren I, Lewis K. (2016). Persisters: Methods for Isolation and Identifying Contributing Factors--A Review. Methods Mol Biol. 1333:17–28.

9. Niumsup P, Simm AM, Nurmahomed K, Walsh TR, Bennett PM, Avison MB. (2003). Genetic linkage of the penicillinase gene, *amp*, and *blrAB*, encoding the regulator of beta-lactamase expression in *Aeromonas* spp. J Antimicrob Chemother. 51:1351–8.

10. Avison MB, Niumsup P, Nurmahomed K, Walsh TR, Bennett PM. (2004). Role of the ‘cre/blr-tag’ DNA sequence in regulation of gene expression by the *Aeromonas hydrophila* beta-lactamase regulator, BlrA. J Antimicrob Chemother. 53:197–202.

11. Tayler AE, Ayala JA, Niumsup P, Westphal K, Baker JA, Zhang L, Walsh TR, Wiedemann B, Bennett PM, Avison MB. (2010). Induction of beta-lactamase production in *Aeromonas hydrophila* is responsive to beta-lactam-mediated changes in peptidoglycan composition. Microbiology. 156:2327–35.

12. Moya B, Dötsch A, Juan C, Blázquez J, Zamorano L, Haussler S, Oliver A. (2009). Beta-lactam resistance response triggered by inactivation of a nonessential penicillin-binding protein. PLoS Pathog. 5:e1000353.

13. Zamorano L, Moyà B, Juan C, Mulet X, Blázquez J, Oliver A. (2014). The *Pseudomonas aeruginosa* CreBC two-component system plays a major role in the response to ß-lactams, fitness, biofilm growth, and global regulation. Antimicrob Agents Chemother. 58:5084–95.

